# DNA-barcoded polysaccharide specific monoclonal antibodies facilitate sensitive and multiplexed detection of cell wall polymers

**DOI:** 10.64898/2026.08.24.746824

**Authors:** C. Frankie Griffith, Michael G. Hahn, Ian S. Wallace

## Abstract

Plant cell walls are polysaccharide-rich extracellular matrices composed of multiple complex carbohydrate polymer networks, including cellulose, hemicelluloses, pectins, and glycosylated proteins. Polysaccharide deposition critically impacts cell wall structure, and structural microheterogeneity within cell wall glycans also influences polymer rigidity and polymer-polymer interactions. Collections of monoclonal antibodies (mAbs) have been developed to target unique carbohydrate epitopes within cell wall polysaccharides and to investigate how these structural changes impact cellular and plant development. Here, we implement generalizable methods to attach unique DNA barcodes to mAbs that recognize major cell wall polysaccharide classes. By applying these mAbs individually to polysaccharide standards, we demonstrate that bound DNA barcoded antibody abundance can be measured via quantitative PCR. Additionally, we demonstrate that DNA conjugated antibodies can be pooled to quantitatively analyze polysaccharide epitope composition of polysaccharide standards and fractionated cell wall material by amplifying their unique barcodes via qPCR. These results demonstrate that barcoded polysaccharide-directed mAbs offer sensitive, quantitative insights into cell wall polysaccharide composition and facilitate multiplexed profiling of cell wall polysaccharide abundance. This approach will also enable multiple future high-throughput applications, such as glycome profiling, spatial glycomics, and glycan interaction measurements, that will further our understanding of cell wall compositional impacts on plant physiology.

## Introduction

Plant cell walls are complex polysaccharide-rich extracellular matrices that surround all plant cells and critically impact basic cellular processes, such as cell expansion, cell division, the acquisition of plant cell shape, and cell adhesion (Cosgrove, 2023; Delmer et al., 2024; Somerville et al., 2004). Plant cell wall glycans can be broadly grouped into four main categories: cellulose, hemicelluloses, pectins, and glycoproteins. However, the composition and fine structure of cell wall polysaccharides can vary dramatically between plant species, within different tissues of the same organism, or in response to environmental perturbations (Burton et al., 2010; Cosgrove, 2023; Delmer et al., 2024; Xiao et al., 2025). It is postulated that cell wall polysaccharide composition and structural microheterogeneity impacts major aspects of cell wall mechanical behavior, but few analytical approaches simultaneously provide high structural and spatial resolution to rigorously investigate this hypothesis.

The substantial structural complexity of plant cell wall polysaccharides has been recently reviewed in detail elsewhere (Cosgrove, 2023; Delmer et al., 2024). Due to this structural complexity, panels of monoclonal antibodies (mAbs) have been developed to recognize specific polysaccharide epitopes within plant cell walls. For example, the Complex Carbohydrate Research Center (CCRC) mAb collection encompasses over 150 mouse-derived mAbs that target carbohydrate epitopes in nearly all major cell wall polysaccharide networks (Pattathil et al., 2010). The John Innes Center Monoclonal (JIM) and Leeds University Monoclonal (LM) series of cell wall antibodies also contain a large number of overlapping and unique probes for the evaluation of plant cell wall polysaccharide structure (Ruprecht et al., 2017). Other laboratories have also generated valuable antibodies that contribute to the size and diversity of the world-wide collection of plant glycan-directed antibodies. mAbs in these collections can recognize very subtle structural differences in plant cell wall polysaccharides, such as xyloglucan fucosylation (Freshour et al., 2003; Puhlmann et al., 1994), 4-O-methylation of glucuronic acid in xylan (Pattathil et al., 2010; Ruprecht et al., 2017; Schmidt et al., 2015), degree of pectin methyl esterification (Clausen et al., 2003; Knox et al., 1990; Verhertbruggen, Marcus, Haeger, Ordaz-Ortiz, et al., 2009; Willats et al., 2001), and polysaccharide degree of polymerization (Ruprecht et al., 2017; Verhertbruggen, Marcus, Haeger, Verhoef, et al., 2009). Current information about the epitopes recognized by plant glycan-directed antibodies has been comprehensively reviewed elsewhere (Thorne et al., 2024). These unique probes have been used for a wide variety of applications, including immunohistochemistry to investigate cell wall polysaccharide distributions in fixed tissues (Verhertbruggen, Marcus, Haeger, Verhoef, et al., 2009), immunochemical profiling of isolated cell wall polysaccharides and their deconstruction products (Biswal et al., 2025; Gunawan et al., 2017; Li et al., 2018), and profiling of mutants with altered cell wall structure (Freshour et al., 2003; Kaur et al., 2022).

Libraries of cell wall mAbs, in conjunction with automated enzyme-linked immunosorbent assays (ELISA), have also been utilized to probe fractionated cell wall polysaccharides to investigate dynamic changes in the plant cell wall glycome in mutants of cell wall metabolism enzymes, during development, or in response to environmental changes. In this approach, total cell wall material (typically alcohol insoluble residue; AIR) is subjected to a series of increasingly basic extractions that progressively solubilize cell wall glycan populations. Fractionated glycans are immobilized in ELISA plates and probed with over 80 different cell wall polysaccharide-directed mAbs to measure quantitative differences in mAb epitope abundance in extracted cell wall fractions. Glycome profiling has provided important insights into cell wall polysaccharide compositional changes in mutants, biomass crops, and even plants exposed to spaceflight (Biswal et al., 2025; De Souza et al., 2015; Nakashima et al., 2023; Okekeogbu et al., 2019; Pattathil et al., 2015; Walker et al., 2017; Wilkop et al., 2019). However, the current implementation of cell wall polysaccharide glycome profiling suffers from a few key limitations, particularly in the context of high-throughput analyses of large sample numbers. First, this approach requires highly specialized liquid handling instrumentation that is inaccessible to most laboratories, thus limiting scope and throughput. Second, all antibodies in the CCRC, JIM, and LM collections were generated in mouse or rat, which limits the number of probes that can be multiplexed in a single reaction due to the species-specific secondary antibody requirements that are necessary to recognize these probes.

In other systems, alternative immunological probe labeling strategies have enabled higher throughput analyses with increased sample multiplexing. For example, attaching specific DNA sequences to antibodies enables antigen abundance to be evaluated by quantitative *in vitro* or spatial approaches (Black et al., 2021; Hoenigsperger et al., 2026; Zhong et al., 2025). Here, we sought to circumvent key limitations associated with traditional glycome profiling by conjugating cell wall polysaccharide-directed mAbs with oligonucleotide barcode tags. We identify a general strategy for installing DNA barcodes onto purified IgG-type cell wall polysaccharide-targeting mAbs and demonstrate that prototypical mAbs considered here can recognize as little as 0.25 ng of target polysaccharide standard when qPCR is used as a detection method. We also demonstrate that barcoded mAbs recognize their cognate polysaccharide targets when applied individually or as a group to pure polysaccharide standards or mixtures of these standards. Lastly, we show that ELISA-based cell wall polysaccharide abundance profiles can be faithfully recreated using this multiplexed method.

## Results

### DNA barcoding of plant cell wall polysaccharide epitope directed antibodies

To initiate this study, we sought a set of mAbs that recognize epitopes within major plant cell wall polysaccharide networks and that exhibited minimal epitope cross reactivity. CCRC-M23, CCRC-M38, CCRC-M58, and CCRC-M140 were initially chosen to meet these criteria because prior work indicates that they selectively recognize unique carbohydrate epitopes in rhamnogalacturonan-I (RG-I), homogalacturonan (HG), xyloglucan (XLG), and arabinoxylan (AX), respectively (Ruprecht et al., 2017). Each of these mAbs also recognize commercially available polysaccharide substrates. To confirm that these mAbs selectively recognized target substrates and exhibited minimal cross reactivity with other polysaccharide standards considered in these initial experiments, replicate samples of soy RG-I (RG-I), apple pectin (Pec), tamarind xyloglucan (XLG), and wheat arabinoxylan (AX) were immobilized into separate wells of a 96-well plate and probed with each antibody using a traditional ELISA-based approach (Supplemental Figure S1). CCRC-M58 exhibited strong ELISA signal against XLG, but was minimally active against Pec, AX, and RG-I substrates. Similarly, CCRC-M140 robustly bound AX, but showed minimal ELISA reactivity against all other substrates tested. CCRC-M38 exhibited the strongest ELISA signal against the Pec polysaccharide standard and CCRC-M23 was most reactive against the Soy RG-I standard. Although the overall ELISA signal was 3 times lower than those observed for CCRC-M58 and CCRC-M140 against their cognate substrates, CCRC-M23 and CCRC-M38 showed their maximum ELISA signals against their cognate substrates and minimal reactivity against other tested polysaccharides. Overall, these results indicate that CCRC-M23, CCRC-M38, CCRC-M58, and CCRC-M140 along with their corresponding polysaccharide standards form a set of reagents that exhibit orthogonal binding activities with minimal cross-reactivity.

Next, we sought a generalized approach for antibody-DNA conjugation to attach specific DNA barcodes for antibody multiplexing. We designed a DNA barcode sequence structure (Figure 1A) that contains an M13 forward primer sequence, a unique 20 nucleotide barcode (BC1), an M13 reverse primer sequence, and a second unique 20 nucleotide barcode (BC2). The rationale for this sequence structure is that the BC1 sequence could be amplified from any barcoded mAb using M13 forward and reverse primers, and that sequences from each antibody could be amplified selectively from a pool using a combination of BC1 and BC2 primers. BC1 and BC2 sequences were chosen from DNA barcodes used to generate the yeast deletion collection barcoded mutant library because these barcode sequences exhibit minimal annealing cross-reactivity (Pierce et al., 2007). To verify that each set of unique barcode sequences could be amplified orthogonally, we synthesized oligonucleotides containing all sequence elements outlined above and used these oligonucleotides as templates for qPCR amplifications with corresponding antisense BC1 and BC2 primer pairs. The resulting qPCR data (Supplemental Figure S2) indicates that each BC1 and BC2 primer pair amplified the target template sequence with minimal crosstalk with other barcode templates. Primer sets targeting CCRC-M23, CCRC-M38, and CCRC-M140 barcodes produced > 100,000 times more amplicon from their target template compared to off target templates. The CCRC-M58 primer set was an exception and exhibited higher background against other barcode sequences, but off-target amplicons were 500-fold less abundant than the on-target barcode sequence. Together, these results demonstrate that these barcode sequences can be selectively amplified by their cognate primer pairs with minimal cross-reactivity against other barcode sequences.

**Figure 1:**
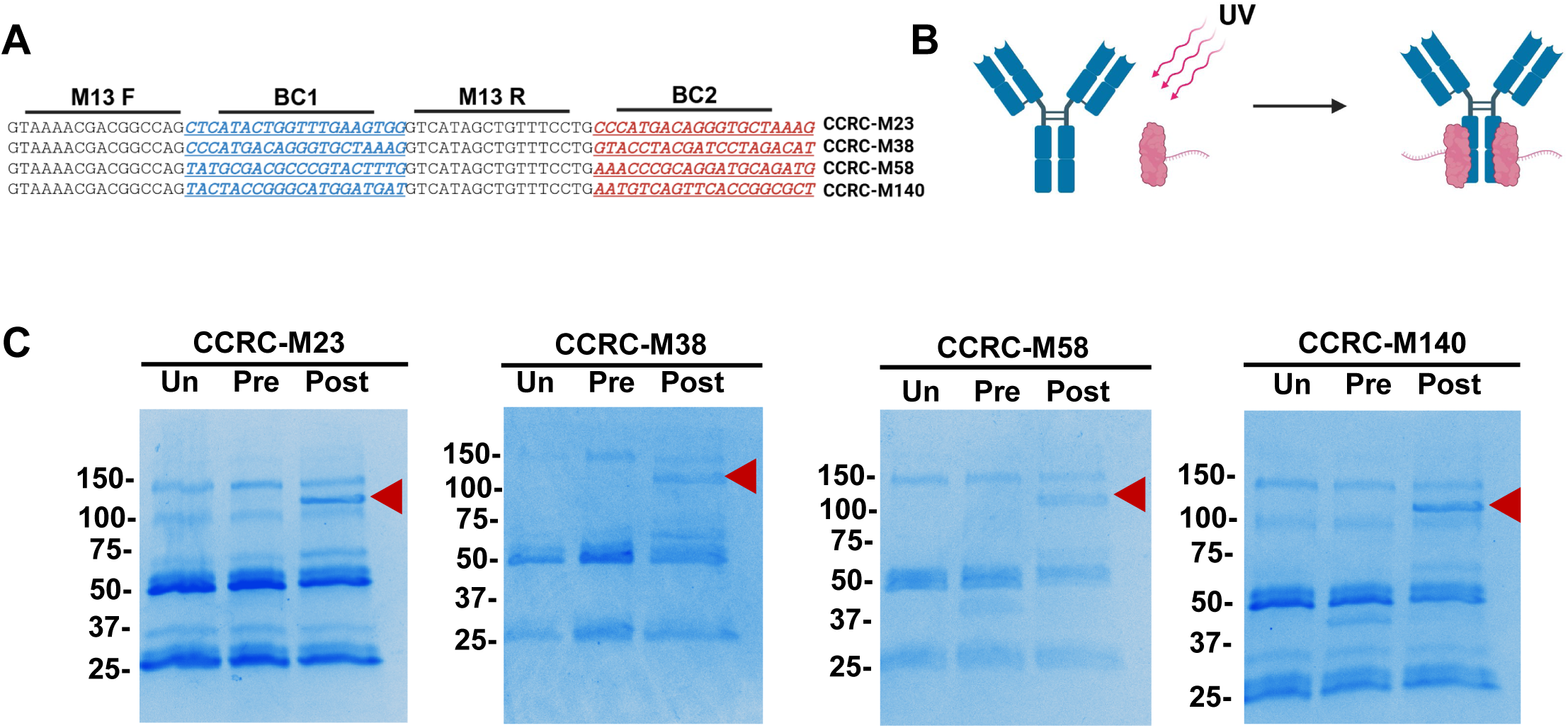
Cell wall mAb DNA conjugation: **A.** The structure of DNA barcodes used in this study are shown. M13 forward and reverse sequences are indicated in black font, while variable BC1 and BC2 sequences are shown in blue and red, respectively. The antibody associated with each barcode is indicated to the right of each sequence. **B.** A schematic of the LASIC labeling process is shown. Protein Z variants containing a photoreactive para benzoyl-L-phenylalanine amino acid in the Fc binding interface are conjugated to a DNA probe (pink). These reagents bind the Fc region of a target antibody (blue) and are specifically crosslinked in the presence of UV light. **C.** The indicated mAb was purified from hybridoma supernatant using Protein G affinity chromatography as described in Materials and Methods. The unmodified antibodies (un) were separated on 4-20% gradient SDS-PAGE gels along with antibody-oYo-Link reagent mixtures before (pre) and after (post) UV irradiation. Red carats indicate the position of putative antibody heavy chain-oYo-Link reagent conjugates. The positions of molecular weight markers are indicated to the left of each gel.

We used Light Assisted Site-Specific Conjugation (LASIC) to attach DNA barcodes to target mAbs (Figure 1B). This approach relies on the light-induced crosslinking of a DNA conjugated protein Z variant containing the photo-crosslinkable amino acid p-benzoyl phenylalanine to the F_c_ region of a target mAb (Hui et al., 2015). LASIC is superior to other antibody conjugation methods, such as introduction of labels through succinimide or maleimide chemistry, because DNA barcode tag installation stoichiometry and regioselectivity is precisely controlled. All target antibodies considered here are mouse IgG_1_ isotypes and were purified from hybridoma supernatant by Protein G chromatography as described in Materials and Methods. IgG_1_-specific LASIC reagents attached to DNA barcode sequences specified in Figure 1A were purchased from AlphaThera. Purified mAbs were incubated with LASIC reagents displaying their cognate DNA barcode and coupled under UV light (365 nm) for 2 hours. LASIC reagent coupling to each antibody was evaluated by SDS-PAGE as compared to the purified, unmodified mAb, and the mAb mixed with LASIC reagent before UV irradiation. As shown in Figure 1C, a heavy chain-LASIC reagent conjugate was observed for each antibody after coupling, signifying that LASIC reagents were successfully attached to each mAb.

Next, we investigated whether DNA barcoded mAbs could quantitatively report carbohydrate antigen content using qPCR as a detection method. To examine this question, soybean RG-I was immobilized in 96 well PCR plates at loading amounts ranging from 100 to 0.25 ng. These wells were probed with DNA-barcoded CCRC-M23 and amplification products from the bound mAbs were quantified by qPCR as described in Materials and Methods. As shown in Figure 2A, a dose-dependent decrease in qPCR amplification and cycle quotient (Cq) was observed when DNA barcoded CCRC-M23 was used to probe increasing soybean RG-I sample loadings, indicating that this probe quantitatively reports immobilized polysaccharide antigens. Plotting the Cq value versus log of antigen loading (Figure 2B) revealed a linear relationship between 10 ng and 0.25 ng of RG-I, suggesting that barcoded mAbs coupled with qPCR as an output sensitively detect target polysaccharides at low loading amounts. Overall, these results suggest that LASIC is a general approach for barcode installation on IgG-type mAbs recognizing plant cell wall polysaccharide epitopes, and that the resulting carbohydrate-specific mAb-DNA conjugates can detect as little as 0.25 ng of their cognate immobilized polysaccharide substrate using qPCR as a detection method.

**Figure 2:**
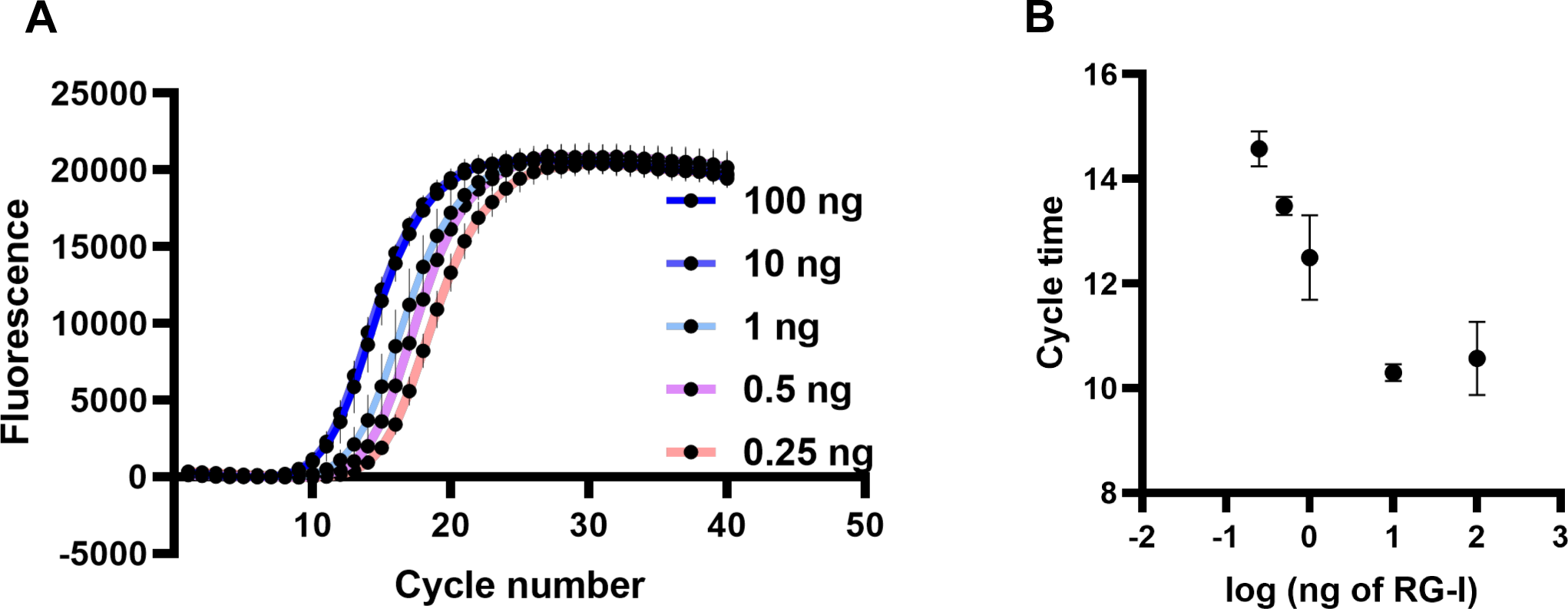
Dose dependence of antigen binding evaluated by immuno-qPCR: Soybean RG-I was immobilized in 96-well PCR plates at quantities ranging from 100 ng to 0.25 ng. Wells were then probed with 0.5 µg/ mL DNA barcoded CCRC-M23 antibody, washed, and subjected to qPCR in the presence of CCRC-M23 barcode-specific primers. **A.** Raw amplification curves from these reactions are shown. **B.** A dose response curve of cycle time as a function of antigen loading is shown. Error bars represent standard deviation (n = 3).

### Evaluating the specificity of individual barcoded antibodies

Next, we evaluated the specificity profile of DNA barcoded CCRC-M23, CCRC-M38, CCRC-M58, and CCRC-M140. Ten nanogram replicate samples of soybean RG-I, citrus pectin, tamarind xyloglucan, and wheat arabinoxylan polysaccharide standards were arrayed into individual wells of a 96-well PCR plate. Additionally, we created an equal mass mixture (Mix) containing 10 ng of each polysaccharide standard that was immobilized separately. Wells with no immobilized polysaccharide (No PS) served as negative controls. These polysaccharide samples were probed with individual DNA barcoded mAbs as described in Materials and Methods. Unbound DNA barcoded mAbs were removed by washing, and barcodes associated with remaining bound antibody were amplified by M13 forward and reverse primers via qPCR (Figure 3A). Amplification curves from these experiments are presented in Supplemental Figure S3A. We used a delta Cq value (dCq) of the maximum cycle time (40 cycles) minus the Cq value of the sample as a quantitative output of amplification (Figure S3B). In contrast to Cq values, dCq value increase with increased amplicon abundance, providing straightforward analysis of positive results. Barcode conjugated CCRC-M23 exhibited the earliest amplification when used to probe its cognate polysaccharide antigen, RG-I. The dCq value of CCRC-M23 applied to RG-I was 27, which was 7-8 cycles earlier than any other individual polysaccharides assayed, representing a > 100-fold increase in amplified product over samples that are not recognized by CCRC-M23. The dCq values of PCR product amplified from the RG-I and mixed polysaccharide samples were 28 and 28.2, respectively. Thus, barcoded CCRC-M23 produced the same amount of amplicon when probed against an individual polysaccharide standard or a defined mixture of polysaccharide standards.

**Figure 3:**
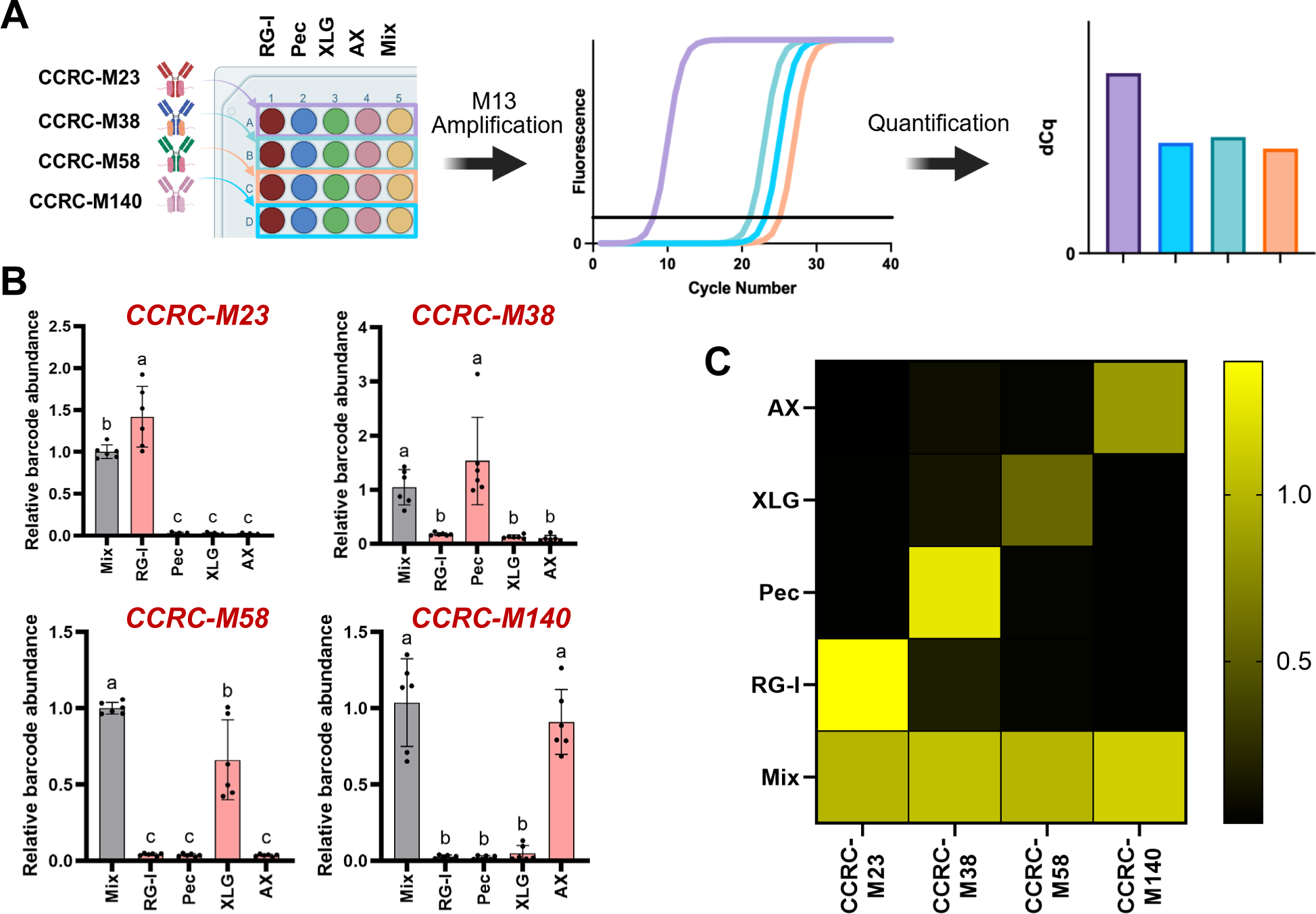
Antigen specificity analysis of barcoded antibodies: **A.** 10 ng of carbohydrate antigens (soybean RG-I (RG-I), citrus pectin (Pec), tamarind xyloglucan (XLG), or wheat arabinoxylan (AX)) or an equal mass mixture (mix) of these polysaccharides were arrayed into wells of a 96-well PCR plate. Each well was probed with barcoded 0.5 µg/ mL DNA barcoded CCRC-M23, CCRC-M38, CCRC-M58, or CCRC-M140 antibodies. Wells containing no polysaccharide (No PS) were included for each antibody as a negative control. After removing unbound antibody, M13 primers were added to each well and bound antibody barcodes were quantified by qPCR. **B.** Amplicon abundance for each sample was calculated relative to the equal mass mixture sample as described in Materials and Methods. Antibodies used to generate each dataset are indicated above the corresponding graph. Error bars represent SD (n = 6; all points shown). Letters above bars represent statistically different categories as evaluated by One-Way ANOVA analysis followed by Tukey’s post-hoc analysis. Samples with unique letters are statistically different from other categories (P<0.001). Samples sharing the same letter are not statistically different from one another. **C.** A confusion matrix heat map of average relative barcode abundance values for all tested samples is shown. A color scale of abundance relative to the mixture sample is indicated to the right of the heatmap.

This pattern was repeated for all barcoded mAbs that were assayed. In all cases, barcoded mAbs amplified with earlier cycle times when applied to their target polysaccharide. dCq values increased by 3-8 cycles, signifying fold changes of 9-256 over non-target antibodies (Supplemental Figure S3). In each case, the average dCq values were not statistically different when a mAb was used to probe an isolated target polysaccharide or the same amount of standard polysaccharide provided in a mixture. Additionally, dCq values for CCRC-M23, CCRC-M38, and CCRC-M140 non-target polysaccharides were not statistically different from negative control reactions containing no polysaccharide. For CCRC-M58, the no polysaccharide control produced a lower dCq value than non-target polysaccharide samples (Figure S3B). To estimate the linear relative abundance differences between these conditions, we subtracted the negative control Cq values from each sample and then calculated ΔΔCq values of each sample from the mixture. We then determined amplicon linear relative abundance by calculating 2^-ΔΔCq^ similarly to standard qPCR gene expression experiments. This approach demonstrated that DNA-barcoded CCRC-M23 produced 100-fold more amplicon when applied to RG-I compared to non-target polysaccharides. The relative abundance of CCRC-M23 amplicons when probing pure RG-I was 1.4-fold higher than the mixture sample, suggesting that this antibody similarly recognizes RG-I in isolation or in mixtures (Figure 3B). This trend was repeated in other antibodies assessed, with CCRC-M38, CCRC-M58, and CCRC-M140 target polysaccharides showing 13.4, 18.3, and 49.5-fold amplicon abundance over non-target polysaccharides and similar relative abundance values between target polysaccharides applied individually or in mixture samples (Figure 3B). These results are summarized in the confusion matrix heatmap of linear relative abundance values presented in Figure 3C. Overall, these results indicate that this group of barcoded mAbs recognize target polysaccharides individually or in a mixed population with amplification selectivity ranging from 13-100 fold.

### Multiplexing of cell wall mAbs for relative quantification using immunoPCR

Next, we investigated whether barcoded mAbs could be applied to polysaccharide samples as a pool and whether bound antibody abundance could be selectively quantified via qPCR using barcode-specific primers (Figure 4A). Barcoded CCRC-M23, CCRC-M38, CCRC-M58, and CCRC-M140 were pooled at equal concentrations of 0.5 µg/ mL and these pooled, DNA conjugated mAbs were applied to PCR plate wells with individual polysaccharide standards, equal mass mixtures, or no polysaccharide negative controls as previously described. After incubation, unbound antibodies were removed by washing, and the abundance of bound mAbs was quantified by qPCR amplification using barcode-specific primers (Figure 4A). Initial amplification curve analysis indicated that primer sets associated with the target antibody-polysaccharide combination yielded earlier amplification than primer sets for non-target polysaccharides for each polysaccharide standard (Supplemental Figure S4). For example, primers targeting the CCRC-M140 DNA barcode amplified 5 cycles earlier when the antibody pool was applied to wells containing AX. In contrast, all other polysaccharides tested with the CCRC-M140 barcode primer set yielded cycle times that were not statistically different from negative control wells. This trend was repeated across all primer sets. In each case, the primer set targeting the barcode for a cognate antibody-polysaccharide combination exhibited dCq values that were higher than all other samples tested, non-target polysaccharides generally exhibited similar dCq values to negative control samples, and the dCq values between individual target polysaccharides and mixtures containing these polysaccharides were nearly identical (Supplemental Figure S4B).

**Figure 4:**
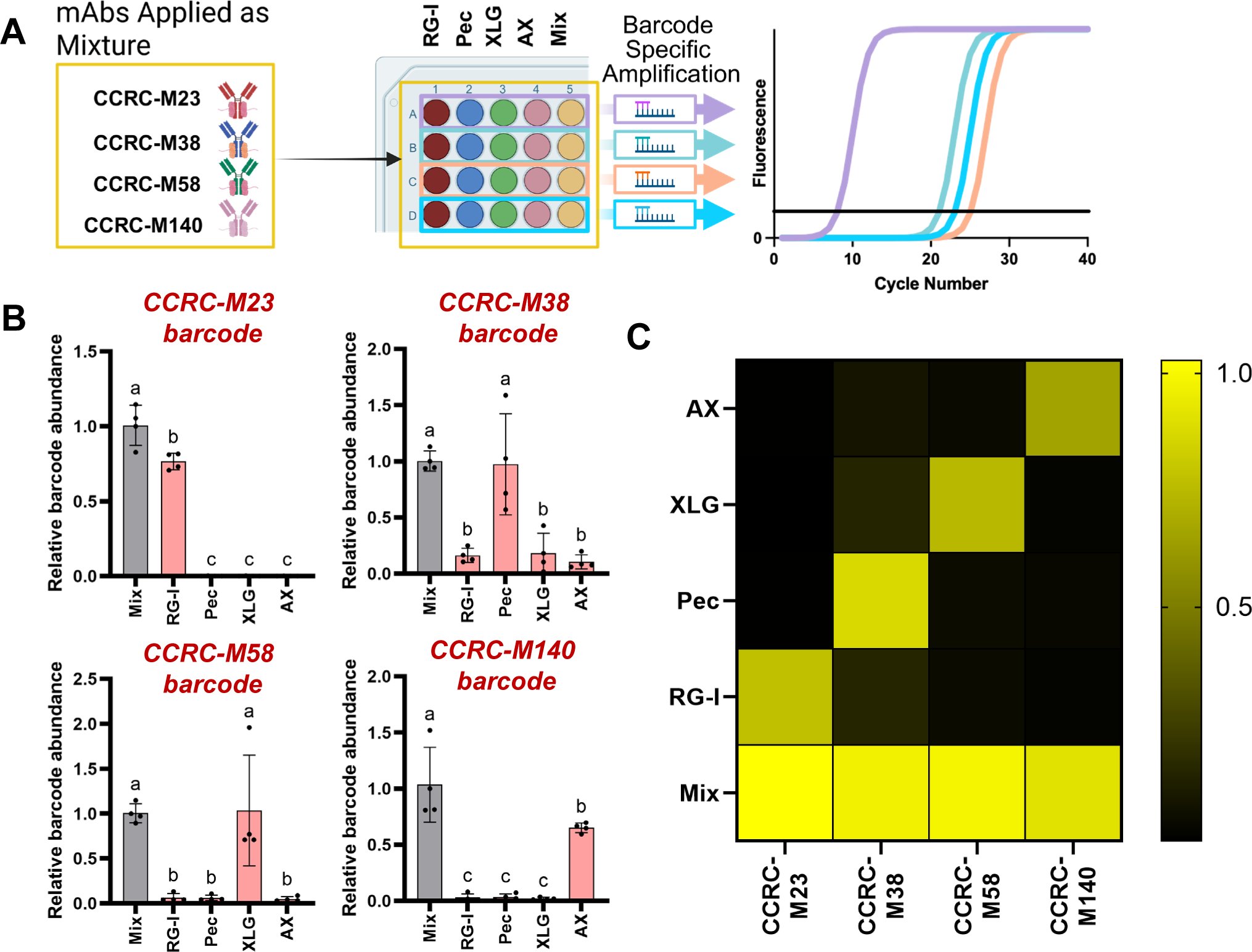
Multiplexed profiling of carbohydrate antigen standards: **A.** 10 ng of carbohydrate antigens were arrayed into replicate wells of a 96-well PCR plate and probed with mixtures of DNA barcoded CCRC-M23, CCRC-M38, CCRC-M58, and CCRC-M140 antibodies each at a concentration of 0.5 µg/ mL. After removing unbound antibody, barcode specific primers were added to each well and bound antibody was quantified by qPCR. **B.** The abundance of each antibody-specific amplicon was calculated relative to the equal mass mixture sample as described in Materials and Methods. Primer sets used to probe the pool of bound antibodies are indicated above each graph. Error bars represent SD (n = 4; all points shown). Letters above bars represent statistically different categories as evaluated by One-way ANOVA analysis followed by Tukey’s post-hoc analysis. Samples labeled with different letters are statistically different (P < 0.001). Samples sharing the same letter are not statistically different from one another. **D.** A heat map of average relative barcode amplicon abundance values for all tested samples is shown. A color scale for the heatmap indicating amplicon abundance relative to the mixture sample is included on the right.

Next, we subtracted negative control Cq values from each sample, generated ΔΔCq values using the polysaccharide mixture as a reference sample and calculated the relative linear abundance of amplicons using the 2^-ΔΔCq^ approach outlined previously. This analysis revealed that CCRC-M23, CCRC-M38, CCRC-M58, and CCRC-M140 associated amplicons were respectively 1634, 11.8, 22, and 40.6-fold more abundant when assayed against target polysaccharides compared to non-target polysaccharides. Additionally, fold changes between target polysaccharides of CCRC-M23, CCRC-M38, CCRC-M58, and CCRC-M140 compared to mixture samples were 0.76, 0.97, 1.03, and 0.62, respectively, suggesting bound antibodies in pure polysaccharide standards or mixture samples could be analyzed by barcode-specific primers with nearly equal efficiency. These results indicate that pooled DNA barcoded cell wall mAbs can quantitatively and selectivity report relative amounts of each mAb-associated epitope in a multiplexed qPCR format when probed with DNA barcode selective primer sets.

### Using barcoded antibodies to quantify fractionated cell wall polysaccharides

Upon verifying that pooled DNA-barcoded mAbs faithfully reported the relative abundance of polysaccharide standards, we sought to understand how this approach performed using fractionated plant biomass samples. Fractionated cell wall polysaccharide samples from developing tomato fruits, Arabidopsis seedlings, and switchgrass stems were obtained, providing a relatively broad compendium of species, tissue types, and cell wall structural features. Switchgrass and tomato samples were fractionated sequentially by ammonium oxalate (AO), sodium carbonate (SC), 1M and 4M KOH. Arabidopsis samples were fractionated by an initial hot water (HW) extraction, followed by AO, 1M and 4M KOH. First, we immobilized 500 ng replicate samples of each polysaccharide fraction in 96 well plates and probed these samples with unmodified CCRC-M23, CCRC-M38, CCRC-M58, and CCRC-M140 using a traditional ELISA approach. The resulting ELISA data (Figure 5A-C) demonstrated antibody reactivity patterns that were consistent with the prior literature (Biswal et al., 2015; Pattathil et al., 2012). For example, pectic epitopes were generally most abundant in the AO and SC extractable polysaccharides for tomato and Arabidopsis samples, but these epitopes were virtually absent in switchgrass samples. CCRC-M58 exhibited higher ELISA signals in the 1M/4M KOH extracts, that typically contain a higher abundance of hemicellulosic epitopes. Lastly, CCRC-M140 ELISA signal was generally low in tomato and Arabidopsis samples but was significantly enriched in the 1M/4M fractions of switchgrass, which is a typical extraction pattern for these tightly bound polysaccharides in each species.

**Figure 5:**
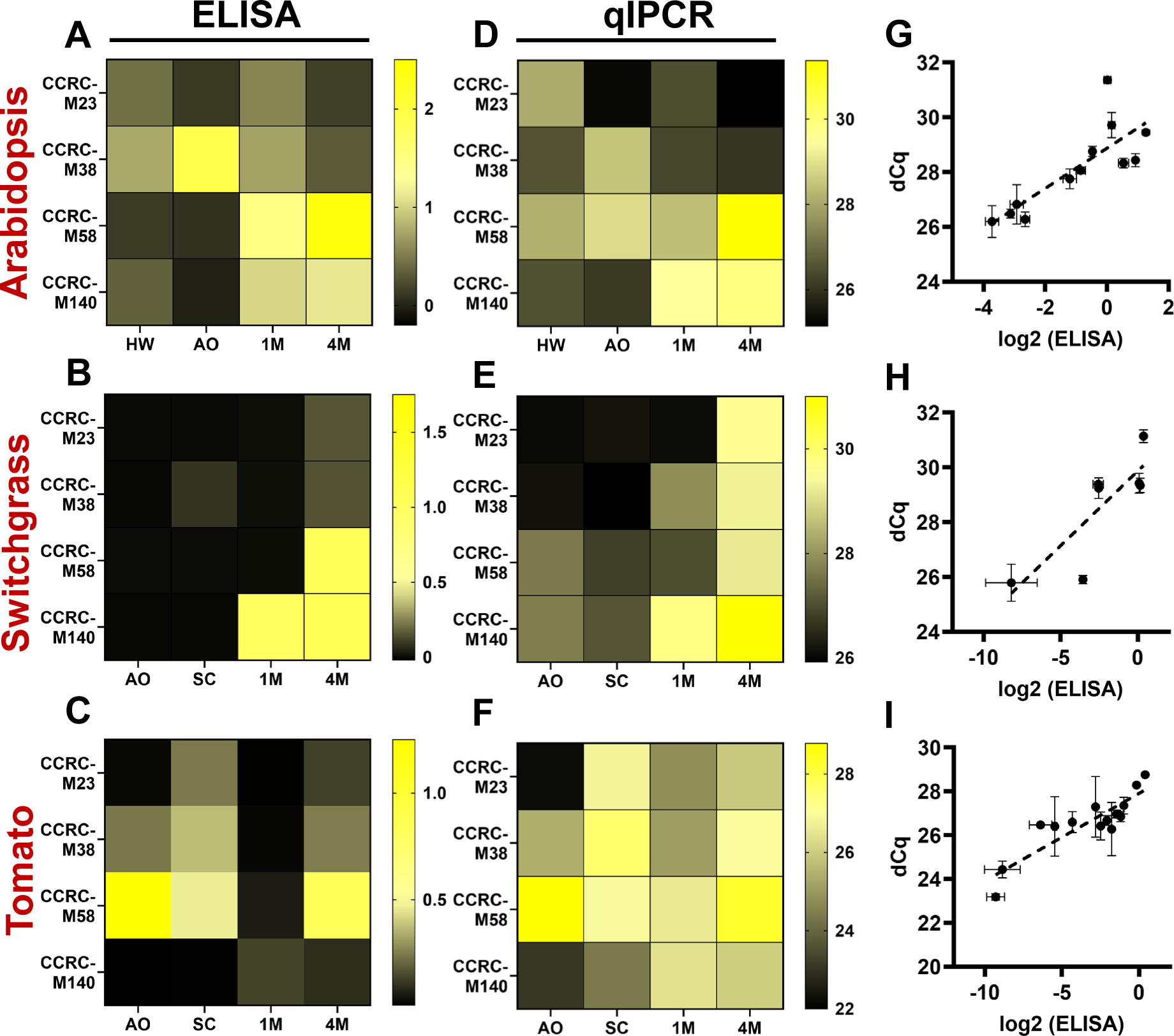
Analysis of fractionated plant cell walls with pooled DNA barcoded antibodies: Arabidopsis, switchgrass, and tomato alcohol insoluble residues were fractionated as described in Materials and Methods. 500 ng of carbohydrates from each fraction were probed with underivatized CCRC-M23, CCRC-M38, CCRC-M58, or CCRC-M140 primary antibodies and subjected to ELISA analysis (A-C). 50 ng of the same samples were immobilized in PCR plates, probed with pools of DNA barcoded mAbs, and bound mAbs were quantified by qIPCR using barcode specific primer sets (D-F). Average A_405_ (A-C) and dCq (D-F) values are represented in each heat map with a corresponding color scale bar to the right of each panel. Fraction abbreviations are hot water (HW), ammonium oxalate (AO), sodium carbonate (SC), 1M KOH (1M), and 4M KOH (4M). dCq values were compared to the log_2_ of the ELISA absorbance signal for Arabidopsis (G), switchgrass (H), and tomato (I) samples. Linear regressions for each dataset are shown as dashed lines. Error bars represent SEM (n = 3).

Next, we immobilized 50 ng replicate samples of these fractions into individual wells of a 96-well PCR plate and probed each fraction with pooled DNA-barcoded mAbs as described above. Barcode-specific primer sets were used to quantify the relative abundance of each mAb by qPCR (Figure 5D-F). Generally, the dCq values for amplicons resulting from antibody-specific primer pairs mirrored ELISA signals of the corresponding antibodies across all fractions. To confirm these trends, we plotted the dCq values against the log_2_ (A_405_) resulting from our ELISA experiments for all samples that produced an ELISA absorbance higher than background (Figure 5G-I). For each group of biomass samples, a linear trend was observed between these parameters, suggesting that quantitative immune-PCR (qIPCR) provided a similar quantitative profile of these biomass fractions as would be expected via traditional ELISA profiling. However, qIPCR identified a variety of low abundance signals, such as CCRC-M23 and CCRC-M38 epitopes in 1M KOH fractions of switchgrass and CCRC-M140 epitopes that were sequentially extracted from tomato cell walls, that were not detected or were poorly detected via ELISA profiling. qIPCR profiling also required 10-fold lower sample input, suggesting that signal sensitivity is increased via this approach. Together, these results suggest that pooled DNA-barcoded mAbs can faithfully reproduce and improve upon ELISA-based glycome profiling results, enabling the rapid quantification of cell wall polysaccharide epitope abundance in plant biomass samples.

## 4. Discussion

Polysaccharide epitope-specific antibodies have enabled numerous analytical approaches in plant cell wall biology, including comprehensive automated compositional analysis by glycome profiling (Nakashima et al., 2023; Pattathil et al., 2015; Walker et al., 2017). However, glycome profiling sample throughput is currently limited by the volume of individual ELISA assays required to generate these glycomics data and access to instrumentation that can complete these tasks, particularly in a high-throughput context with large sample numbers. Here, we utilized a regioselective approach to install unique DNA barcodes onto plant cell wall polysaccharide-directed mAbs (Figure 1). We then demonstrated that these DNA barcoded mAbs recognize antigens at loading amounts as low as 0.25 ng when using qPCR as a detection method for mAb binding (Figure 2). Subsequently, we showed that a collection of four mAbs that target epitopes unique to different cell wall polysaccharides could recognize their cognate antigens in isolation, in mixtures of polysaccharides, and in multiplexed mixtures of barcoded mAbs (Figures 3 and 4). Amplification of barcode-specific sequences allowed for sensitive detection of specific antibodies in pooled samples (Figure 4). Lastly, we used this pooled mAb approach to profile fractionated cell wall samples from various plant species and tissues, demonstrating that multiplexed barcoded mAbs can recreate cell wall epitope profiles generated by a traditional ELISA-based approach at 10-fold reduced sample loadings, while simultaneously detecting low abundance carbohydrate epitope features that are difficult to observe via ELISA (Figure 5). Overall, these results demonstrate that DNA-barcoded polysaccharide-specific antibodies can be employed as useful and sensitive probes to investigate plant cell wall polysaccharide compositions.

The conjugation of DNA fragments to immunological probes has recently revolutionized several traditional immunological approaches. For example, the spatial proteomics technique Co-Detection by indexing (CODEX) relies on DNA-barcoded antibodies to facilitate high dimensional multiplexed imaging of greater than 40 probes against a single tissue section (Black et al., 2021; Goltsev & Nolan, 2023). This process is enabled by cyclically flowing fluorescently labeled antisense oligonucleotides across the tissue section, which bind to their cognate barcoded mAb. This process does not rely on secondary antibodies and therefore the number of multiplexed probes could be increased substantially provided that the DNA probes used for detection remain orthogonally specific. Additionally, proximity-ligation strategies enable the *in vitro* or spatial detection of adjacent protein epitopes, and this technique is also enabled by DNA-tagged antibodies (Fredriksson et al., 2002; Hoenigsperger et al., 2026). In this approach, antibodies recognizing different antigens that are predicted to be in close proximity are attached to DNA sequences that share partial base pair complementarity. If the antigens are within 40 nm of one another, the DNA strands from DNA-tagged antibodies anneal, enabling amplification techniques, such as standard PCR or rolling circle amplification, to facilitate the detection of these spatial interactions. Finally, DNA barcoded mAbs or nanobodies have been utilized in a similar manner as described here for multiplexed quantification of protein antigens, but short read sequencing is used as a detection method for the high-content quantification of many barcodes in a single sample (Zhong et al., 2025). Thus, the work reported here lays a strong foundation for pursuing these advanced, high-throughput, and information-rich methods to investigate plant cell wall carbohydrate-focused questions. Recently, similar approaches were utilized in mammalian systems to facilitate multiplexed quantification of N- and O-proteoglycans as well as single cell sequencing of cells that exhibit disrupted glycosylation of these epitopes, suggesting that similar approaches will be possible in plant systems (Marglous et al., 2025).

Conceivably, this DNA sequence-facilitated multiplexing approach could be extended to other types of carbohydrate-binding probes to examine the dynamics of extracellular matrices of plants and other organisms. Carbohydrate Binding Modules (CBMs), lectins, and certain extracellular proteins (e.g. mammalian Fibroblast Growth Factors, antithrombins) bind to polysaccharides with high specificity, affinity, and selectivity. Thus, the installation of DNA barcodes on these select probes could increase the scope of polysaccharides profiled using this approach. While traditional labeling chemistries could facilitate this approach, protein ligation methods, such as sortase or asparaginyl endopeptidase-mediated ligation (Dorr et al., 2014; Hemu et al., 2019), may provide increased specificity, regioselectivity, and flexibility for probe installation. Overall, the results presented here lay a strong foundation to enable these high-throughput approaches for glycan characterization in a wide range of biological systems.

## Materials and Methods

### Antibody purification

Hybridoma supernatants containing CCRC-M23, CCRC-M38, CCRC-M58, and CCRC-M140 antibodies were obtained from CarboSource (Athens, GA). Two hundred microliters of NAb protein G resin slurry (NAb Protein G Spin Column; ThermoFisher Scientific) was resuspended in 400 µL of binding buffer (100 mM sodium phosphate pH 7.2, 150 mM NaCl) and transferred to 10 mL of hybridoma supernatant. The resin was incubated with hybridoma supernatant for 2 hours on a nutating table at 25° C. After incubation, the resin was centrifuged at 1,000 x *g* for 1 minute, and the supernatant was decanted. The resin was resuspended in 400 µL of binding buffer and transferred to the NAb spin column. The column was centrifuged at 5,000 x *g* for 1 minute. The supernatant was removed, and the resin was washed 5 times with 400 µL of binding buffer with centrifugation at 5,000 x *g* for 1 minute between each wash. Purified mAbs were eluted by applying 400 µL of Elution Buffer (100 mM glycine pH 2.5), centrifuging the resin, and transferring the supernatant into 40 µL of Neutralization Buffer (1 M sodium phosphate pH 8.5). This process was repeated 5 times. Antibody purity was assessed by SDS-PAGE and protein concentrations were determined by BCA assay (ThermoFisher Scientific). Fractions containing affinity purified mAbs were concentrated using a Pierce 10 KDa Molecular Weight Cutoff (MWCO) protein concentrator (ThermoFisher Scientific). Antibodies were then buffer exchanged with 5 column volumes of binding buffer and concentrated to a final volume of 100 µL. Purified mAbs were adjusted to a final concentration of 0.5 mg/ mL with binding buffer.

### LASIC crosslinking of DNA barcodes

DNA conjugated LASIC reagents covalently attached to DNA barcodes specific for each polysaccharide epitope-directed mAb were custom synthesized by AlphaThera (Philadelphia, PA). Unique single stranded oligonucleotide sequences for CCRC-M23, M38, M58, and M140 (Supplemental Table I) were attached to LASIC reagents during synthesis. Each reagent was suspended in 25 µL of sterile dH_2_O and combined with 75 µL of its corresponding antibody at a final concentration of 0.4 mg/ mL in binding buffer. UV-induced crosslinking was carried out using a LED PX device (AlphaThera) by irradiating the samples at 365 nm for 2 hours at 4° C. Crosslinking was validated by SDS-PAGE.

### ELISA methods

Tamarind xyloglucan, soybean rhamnogalacturonan-I, wheat arabinoxylan (Megazyme), and apple pectin (Sigma Aldrich) were dissolved in sterile water at a concentration of 2 ng/ µL. Total carbohydrates in biomass fractions were quantified by phenol sulfuric acid assay, and these fractions were adjusted to a concentration of 10 ng sugar equivalent/ µL. In a high binding clear 96-well plate (ThermoFisher Scientific), 50 µL of polysaccharide standard or biomass fraction was dispensed into wells and dried overnight at 40° C. Two hundred microliters of Blocking Buffer (1% [w/v] non-fat dry milk (NFDM), 10 mM Tris-HCl pH 7.6, 150 mM NaCl) was added to each well and incubated at 25° C for 1 hour. The buffer was removed and each well was incubated for 1 hour at 25° C with 50 µL of primary antibody in a 1:10 dilution in 0.1% [w/v] NFDM in TBS. The primary antibody solution was removed from each well, and wells were then washed 3 times with 300 µL of Wash Buffer (0.1% [w/v] NFDM, 10 mM Tris-HCl pH 7.6, 150 mM NaCl). The remaining buffer was removed from each well, and 50 µL of secondary antibody solution (1:5000 anti-mouse IgG alkaline phosphatase conjugate [Novus] in wash buffer) was added to each well. The plate was incubated at 25° C for 1 hour, the antibody solution was removed, and each well was washed 3 times with Wash Buffer. After removing all remaining wash buffer, a 5 mg PNPP substrate tablet (ThermoFisher Scientifc) was dissolved in 5 mL of 1X Pierce diethanolamine PNPP substrate buffer (ThermoFisher Scientific). One hundred microliters of substrate was added to each well and incubated at 25° C for 30 minutes in the dark. The reactions were terminated by the addition of 50 µL of 2 N NaOH, and the absorbance of each well was measured at 405 nm using an Agilent BioTek HT multimodal plate reader.

### Quantitative immuno-PCR

Polysaccharides standards described above were dissolved in sterile water at a concentration of 0.2 ng/ µL. Additionally, these four polysaccharides were combined 1:1:1:1 at 0.2 ng/ µL each (0.8ng/ µL total). Biomass fractions were adjusted to a standard concentration of 1 ng/ µL. Fifty microliters of polysaccharide standards were arrayed into individual wells of 96-well polypropylene PCR plates (Fisher Scientific) and dried for 24 hours in a heating block at 40° C. Each well was blocked with 200 µL of Blocking Buffer for 1 hour at 25° C. The blocking buffer was removed, and 50 µL of antibody-DNA conjugate at a concentration of 0.5 ng/µL diluted in Wash Buffer was added. For experiments that required antibodies to be applied as a pool, 0.5 ng/ µL of each barcoded antibody was applied in the mixture. After incubation for 1 hour at 25° C, the antibody solution was removed, and each well was washed 3 times with 200 µL of Wash Buffer. The remaining buffer was removed by blotting onto paper towels, and 10 µL of 1X Sso Advanced SYBR Green Universal Supermix (Bio-Rad) containing 500 nM primers (Supplemental Table II) was added to each well. Samples were analyzed by quantitative PCR using an Agilent mx3000p thermocycler under the following cycling conditions: initial denaturation at 95° C for 30 seconds, 40 cycles of 95° C for 10 seconds followed by 55° C for 10 seconds and fluorescence measurement. Melting curves were also constructed after the final amplification cycle by ramping the temperature from 65° C to 95° C in 0.5° C increments with a fluorescence measurement at each temperature increment.

### Fractionated Cell Wall Preparations

Alcohol insoluble residue was prepared from 1 g of Arabidopsis Col-0 seedlings, *Solanum pimpinellifolium* LA1589 fruits, or Panicum virgatum cv. Alamo stems as previously described (Biswal et al., 2018; Villalobos et al., 2015). Carbohydrate fractionation was conducted as previously described (Pattathil et al., 2012). Total polysaccharide content was estimated using the phenol sulfuric acid assay (Pattathil et al., 2012). Fractions were adjusted to 10 ng/ µL for ELISA profiling and 1 ng/ µL for qIPCR profiling.

## Acknowledgements

We would like to thank our Complex Carbohydrate Research Center colleagues Jason Backe, Breeanna Urbanowicz, Ajay Biswal, and Debra Mohnen as well as Yanbing Wang and Esther van der Knapp (University of Georgia Department of Horticulture) for providing fractionated biomass samples.

## Data Availability Statement

All data necessary to evaluate the conclusions of this study are included in the manuscript and supplementary information. Raw datasets and additional research materials are available from the corresponding author upon reasonable request.

---

**Supplemental Figure S1:**
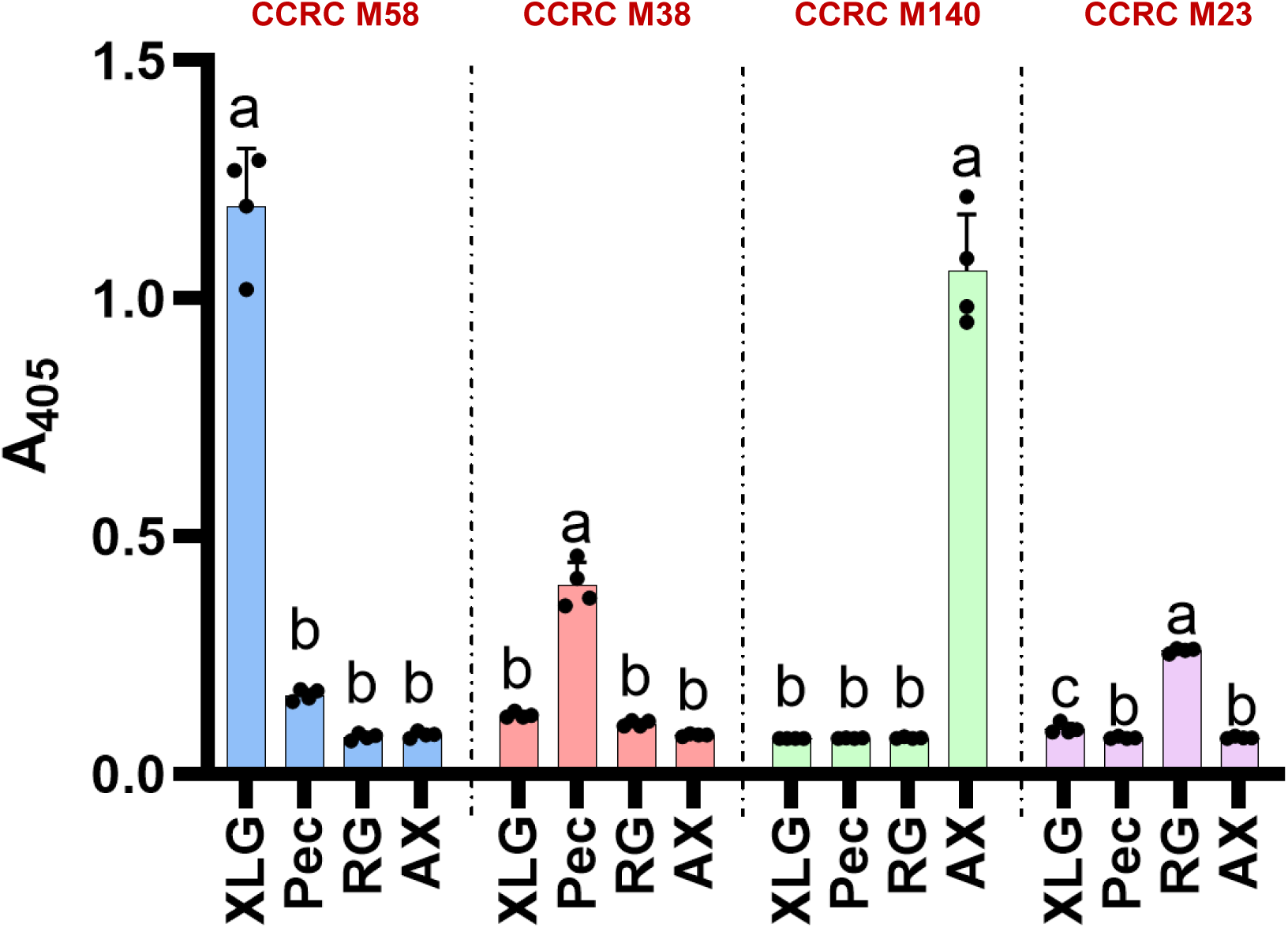
ELISA-based cross-reactivity assays of CCRC mAbs considered in this study: 96 well plates were coated with 100 ng of Tamarind Xyloglucan (XLG), Apple pectin (Pec), Soybean RG-I (RG), or Wheat Arabinoxylan (AX). Wells were blocked and probed with 1:10 dilutions of CCRC-M58 (blue bars), CCRC-M38 (red bars), CCRC-M140 (green bars), or CCRC-M23 (magenta bars) hybridoma supernatant. ELISA signals were generated by a goat anti-mouse alkaline phosphatase conjugated secondary antibody as described in Materials and Methods. Data are plotted with all data points shown. Error bars represent SD (n = 4). Letters above bars represent statistically different categories as determined by One-Way ANOVA analysis of individual antibody groups followed by Tukey’s post-hoc analysis (P<0.001). Samples sharing the same letter are not statistically different from one another.

**Supplemental Figure S2:**
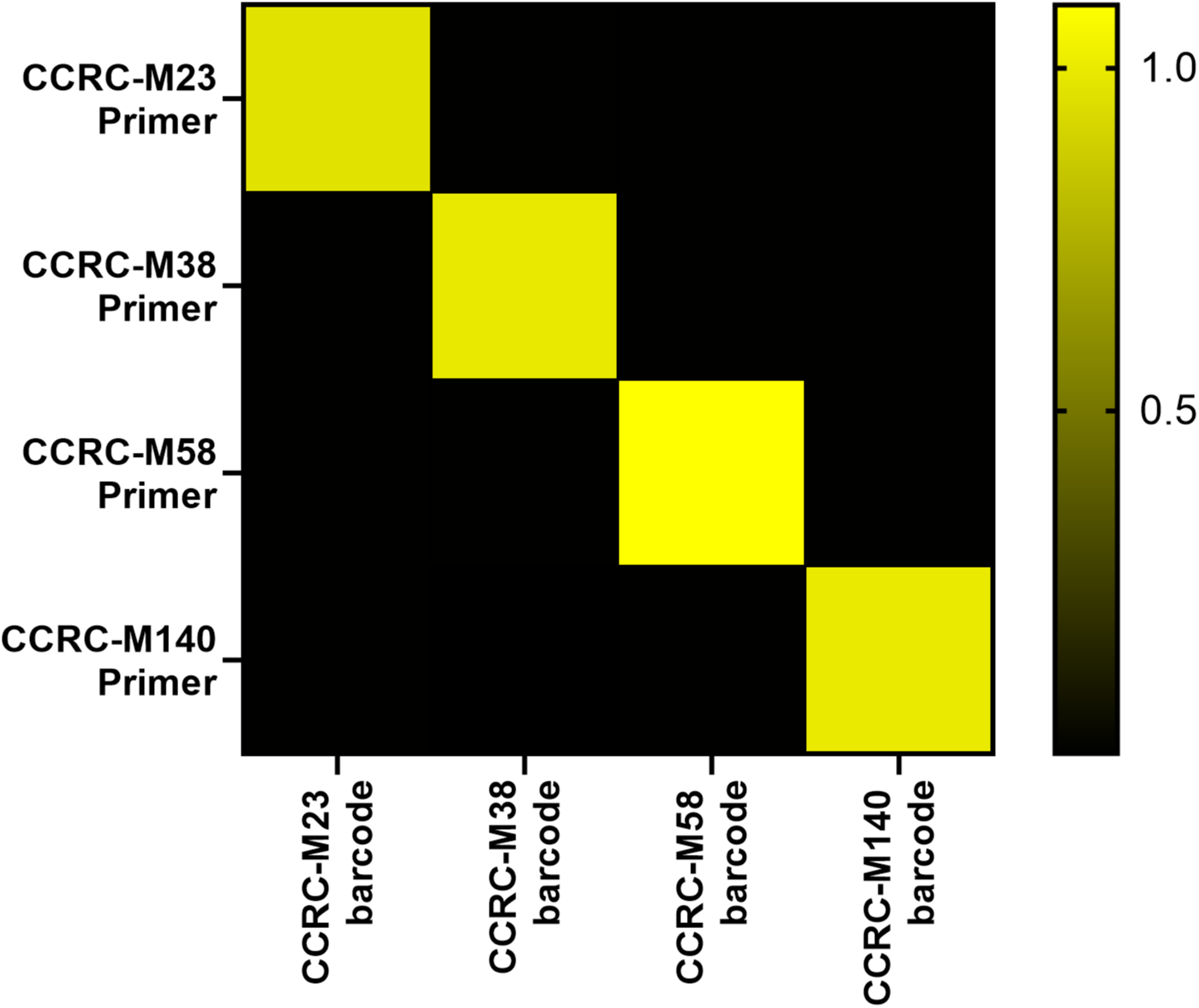
Barcode primer set amplification specificity: Primer pairs (500 nM per primer) specific for the barcode sequences of CCRC-M23, CCRC-M38, CCRC-M58, and CCRC-M140 were incubated with 2 pg of isolated DNA barcode sequence under standard qPCR conditions as described in Materials and Methods. A heat map is shown of the linear amplicon abundance values relative to the target barcode sequence resulting from an all-by-all comparison of these amplification trials. Relative abundance values are colored according to the scale shown on the right. Average relative abundance values are shown (n = 3).

**Supplemental Figure S3.**
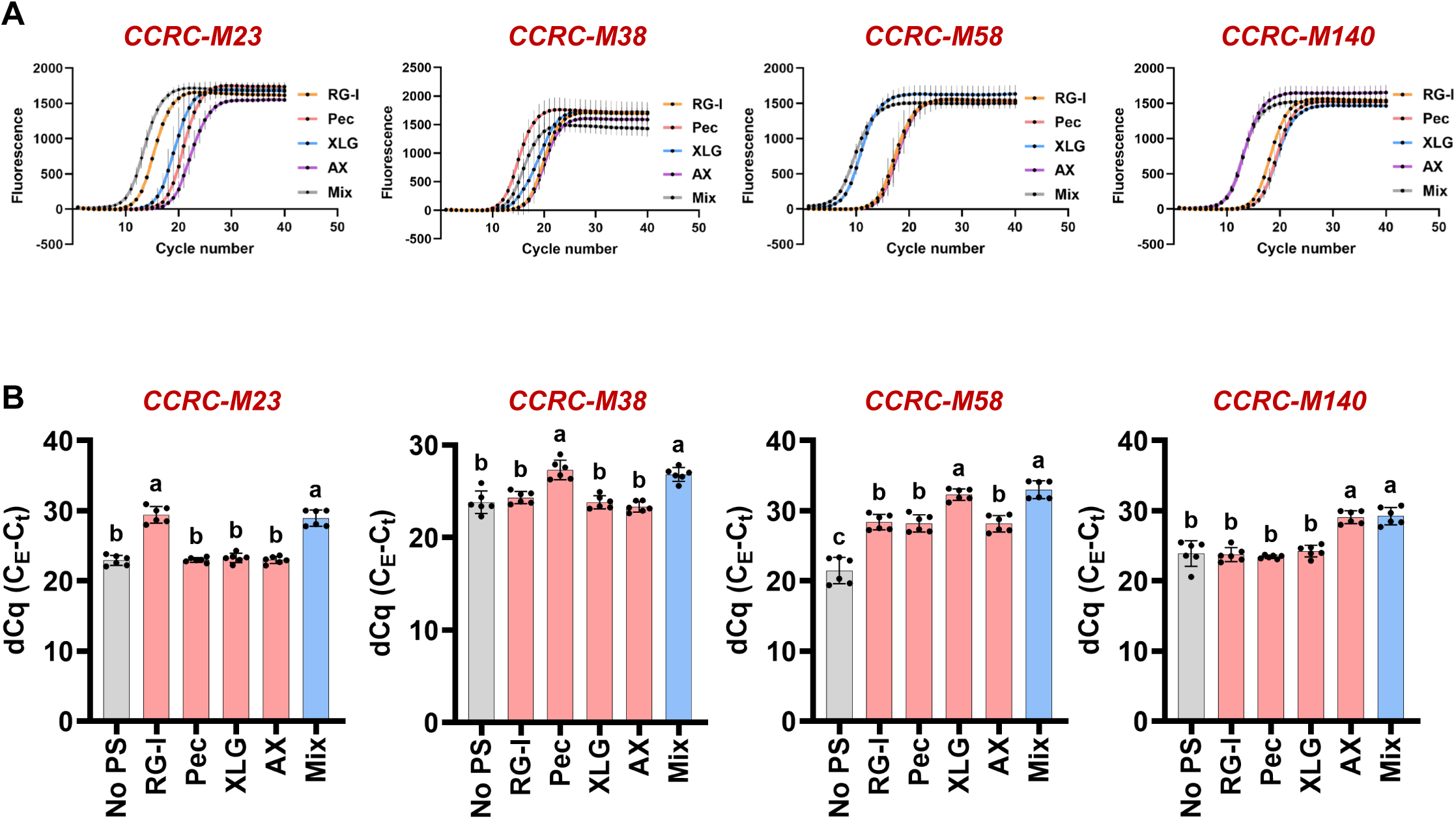
Amplification curves and cycle times of antibody selectivity experiment. **A.** Representative amplification curves are shown for the indicated antibody applied to soy RG-I (RG-I; orange line), apple pectin (Pec; red line), tamarind xyloglucan (XLG; blue line), wheat arabinoxylan (AX; purple line), or an equal part mixture of these polysaccharides (Mix; gray line). Error bars represent SD (n = 3). **B.** The dCq values (Max cycle-Cq value) extracted from amplification curves resulting from each polysaccharide being probed by the indicated antibody are shown. Error bars represent SD (n = 6; all points shown). Letters above bars represent statistically different categories as evaluated by One-Way ANOVA analysis followed by Tukey’s post-hoc analysis. Letters indicate sample categories that are statistically different from one another (P<0.0001). Samples sharing the same letter are not statistically different from each another.

**Supplemental Figure S4:**
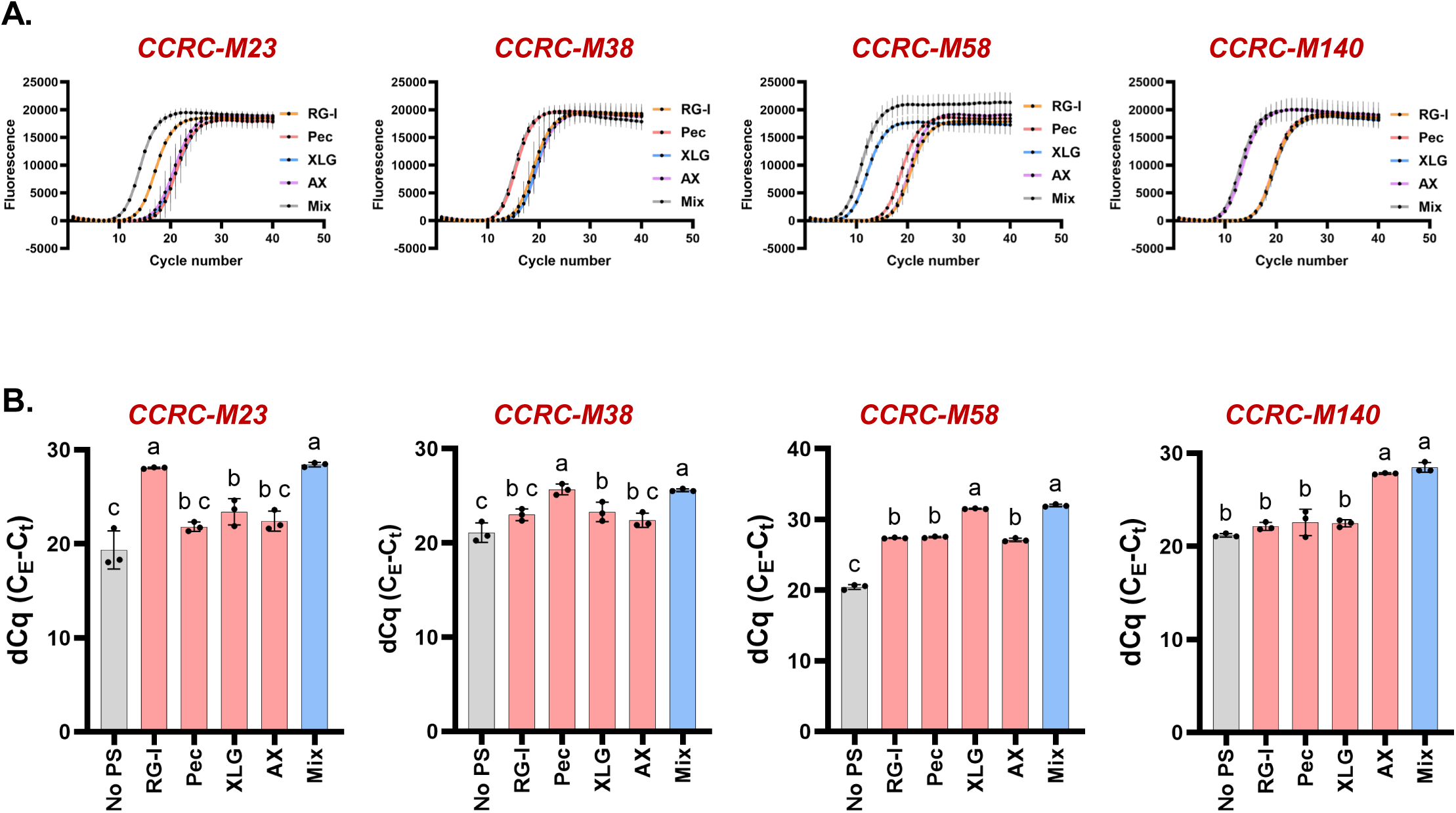
Antigen specificity analysis of barcoded antibodies: **A.** 10 ng of carbohydrate antigens or an equal mass mixture (Mix) were arrayed replicate wells of a 96-well PCR plate and then probed with pooled DNA barcoded antibodies. After removing unbound antibody, barcode-specific primers were added to each well and bound antibody barcodes were quantified by qPCR. **B.** Representative amplification curves are shown for the indicated barcode primer set applied to soy RG-I (RG-I; orange line), apple pectin (Pec; red line), tamarind xyloglucan (XLG; blue line), wheat arabinoxylan (AX; purple line), or an equal part mixture of these polysaccharides (Mix; gray line). Error bars represent SD (n = 3). **C.** The dCq values (Max cycle-Cq value) extracted from amplification curves resulting from each polysaccharide being probed by the indicated antibody are shown. A no polysaccharide (No PS) negative control was included in each experiment. Error bars represent SD (n = 3; all points shown). Letters above bars represent statistically different categories as evaluated by One-Way ANOVA analysis followed by Tukey’s post-hoc analysis. Samples labeled with different letters are statistically different from one another (P<0.001). Samples sharing the same letter are not statistically different from one another.

**Supplemental Table I:**
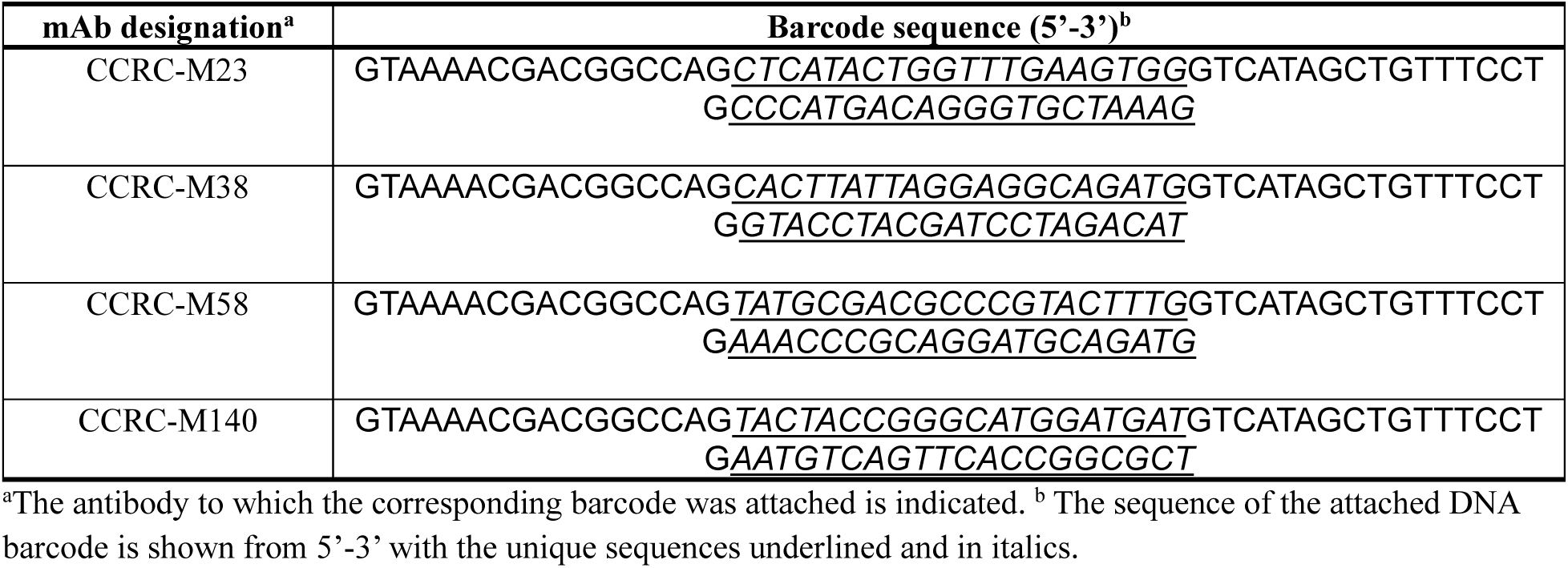
DNA barcode sequences used in this study.

**Supplemental Table II:**
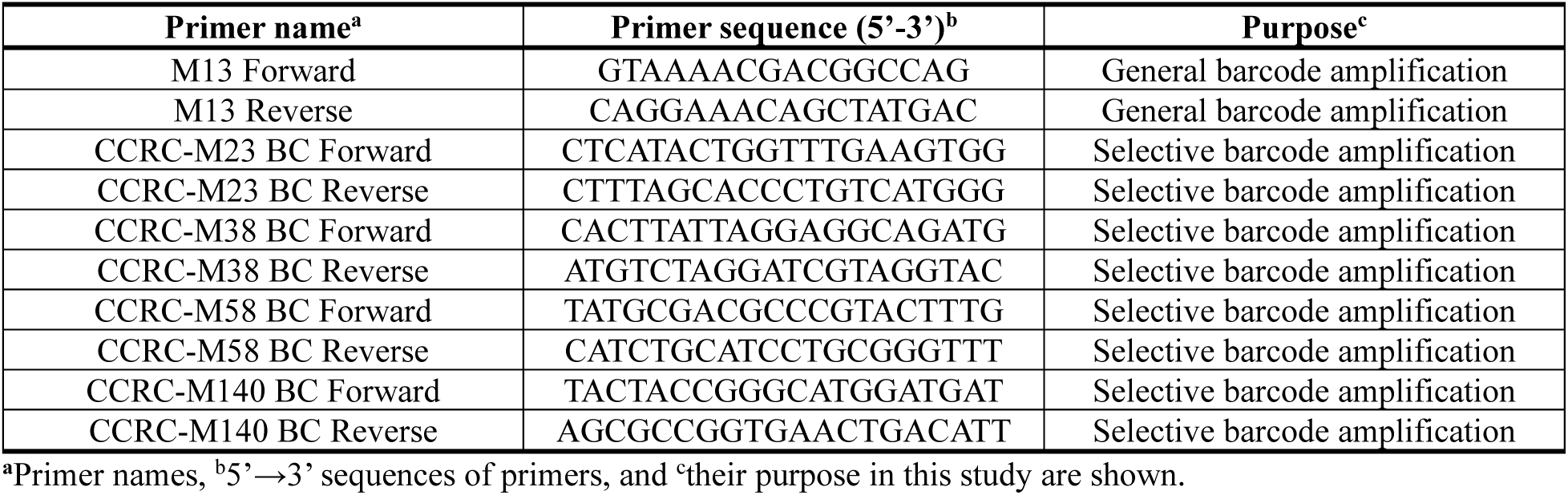
Oligonucleotide primers used in this study.

## References

Biswal, A. K., Atmodjo, M. A., Li, M., Baxter, H. L., Yoo, C. G., Pu, Y., Lee, Y.-C., Mazarei, M., Black, I. M., Zhang, J.-Y., Ramanna, H., Bray, A. L., King, Z. R., Lafayette, P. R., Pattathil, S., Donohoe, B. S., Mohanty, S. S., Ryno, D., Yee, K.,…Mohnen, D. (2018). Sugar release and growth of biofuel crops are improved by downregulation of pectin biosynthesis. Nature Biotechnology, 36(3), 249–257. 10.1038/nbt.4067

Biswal, A. K., Hao, Z., Pattathil, S., Yang, X., Winkeler, K., Collins, C., Mohanty, S. S., Richardson, E. A., Gelineo-Albersheim, I., Hunt, K., Ryno, D., Sykes, R. W., Turner, G. B., Ziebell, A., Gjersing, E., Lukowitz, W., Davis, M. F., Decker, S. R., Hahn, M. G., & Mohnen, D. (2015). Downregulation of GAUT12 in Populus deltoides by RNA silencing results in reduced recalcitrance, increased growth and reduced xylan and pectin in a woody biofuel feedstock. Biotechnology for Biofuels, 8(1), 41. 10.1186/s13068-015-0218-y

Biswal, A. K., Hengge, N. N., Atmodjo, M. A., Abraham, P. E., Engle, N. L., Mohanty, S. S., Black, I. M., Cheng, X., Ryno, D., Azadi, P., Tschaplinski, T. J., Bomble, Y. J., & Mohnen, D. (2025). Rhamnogalacturonan I is a recalcitrant pectin domain during Clostridium thermocellum-mediated deconstruction of switchgrass biomass. Biotechnology for Biofuels and Bioproducts, 18(1). 10.1186/s13068-025-02703-9

Black, S., Phillips, D., Hickey, J. W., Kennedy-Darling, J., Venkataraaman, V. G., Samusik, N., Goltsev, Y., Schürch, C. M., & Nolan, G. P. (2021). CODEX multiplexed tissue imaging with DNA-conjugated antibodies. Nature Protocols, 16(8), 3802–3835. 10.1038/s41596-021-00556-8

Burton, R. A., Gidley, M. J., & Fincher, G. B. (2010). Heterogeneity in the chemistry, structure and function of plant cell walls. Nature Chemical Biology, 6(10), 724–732. 10.1038/nchembio.439

Clausen, M. H., Willats, W. G. T., & Knox, J. P. (2003). Synthetic methyl hexagalacturonate hapten inhibitors of anti-homogalacturonan monoclonal antibodies LM7, JIM5 and JIM7. Carbohydrate Research, 338(17), 1797–1800. 10.1016/S0008-6215(03)00272-6

Cosgrove, D. J. (2023). Structure and growth of plant cell walls. Nature Reviews Molecular Cell Biology. 10.1038/s41580-023-00691-y

De Souza, A. P., Kamei, C. L. A., Torres, A. F., Pattathil, S., Hahn, M. G., Trindade, L. M., & Buckeridge, M. S. (2015). How cell wall complexity influences saccharification efficiency in *Miscanthus sinensis*. Journal of Experimental Botany, 66(14), 4351–4365. 10.1093/jxb/erv183

Delmer, D., Dixon, R. A., Keegstra, K., & Mohnen, D. (2024). The plant cell wall—dynamic, strong, and adaptable—is a natural shapeshifter. The Plant Cell, 36(5), 1257–1311. 10.1093/plcell/koad325

Dorr, B. M., Ham, H. O., An, C., Chaikof, E. L., & Liu, D. R. (2014). Reprogramming the specificity of sortase enzymes. Proceedings of the National Academy of Sciences, 111(37), 13343–13348. 10.1073/pnas.1411179111

Fredriksson, S., Gullberg, M., Jarvius, J., Olsson, C., Pietras, K., Gústafsdóttir, S. M., Östman, A., & Landegren, U. (2002). Protein detection using proximity-dependent DNA ligation assays. Nature Biotechnology, 20(5), 473–477. 10.1038/nbt0502-473

Freshour, G., Bonin, C. P., Reiter, W.-D., Albersheim, P., Darvill, A. G., & Hahn, M. G. (2003). Distribution of Fucose-Containing Xyloglucans in Cell Walls of the mur1 Mutant of Arabidopsis. Plant Physiology, 131(4), 1602–1612. 10.1104/pp.102.016444

Goltsev, Y., & Nolan, G. (2023). CODEX multiplexed tissue imaging. Nature Reviews Immunology, 23(10), 613–613. 10.1038/s41577-023-00936-z

Gunawan, C., Xue, S., Pattathil, S., Da Costa Sousa, L., Dale, B. E., & Balan, V. (2017). Comprehensive characterization of non-cellulosic recalcitrant cell wall carbohydrates in unhydrolyzed solids from AFEX-pretreated corn stover. Biotechnology for Biofuels, 10(1). 10.1186/s13068-017-0757-5

Hemu, X., El Sahili, A., Hu, S., Wong, K., Chen, Y., Wong, Y. H., Zhang, X., Serra, A., Goh, B. C., Darwis, D. A., Chen, M. W., Sze, S. K., Liu, C.-F., Lescar, J., & Tam, J. P. (2019). Structural determinants for peptide-bond formation by asparaginyl ligases. Proceedings of the National Academy of Sciences, 116(24), 11737–11746. 10.1073/pnas.1818568116

Hoenigsperger, H., Klute, S., Engels, Z., Volcic, M., & Sparrer, K. M. J. (2026). Protocol to differentially quantify spatially resolved viral protein-cellular protein interactions via proximity ligation assays. STAR Protocols, 7(1), 104361. 10.1016/j.xpro.2026.104361

Hui, J. Z., Tamsen, S., Song, Y., & Tsourkas, A. (2015). LASIC: Light Activated Site-Specific Conjugation of Native IgGs. Bioconjugate Chemistry, 26(8), 1456–1460. 10.1021/acs.bioconjchem.5b00275

Kaur, D., Moreira, D., Coimbra, S., & Showalter, A. M. (2022). Hydroxyproline-O-Galactosyltransferases Synthesizing Type II Arabinogalactans Are Essential for Male Gametophytic Development in Arabidopsis. Frontiers in Plant Science, 13. 10.3389/fpls.2022.935413

Knox, J. P., Linstead, P., King, J., Cooper, C., & Roberts, K. (1990). Pectin esterification is spatially regulated both within cell walls and between developing tissues of root apices. Planta, 181(4). 10.1007/bf00193004

Li, Y., Zhuo, J., Liu, P., Chen, P., Hu, H., Wang, Y., Zhou, S., Tu, Y., Peng, L., & Wang, Y. (2018). Distinct wall polymer deconstruction for high biomass digestibility under chemical pretreatment in Miscanthus and rice. Carbohydrate Polymers, 192, 273–281. 10.1016/j.carbpol.2018.03.013

Marglous, S., Bhakta, A. Z., Gillmann, K. M., Temme, J. S., Yamamoto, N., Tytla, A., Bahnick, A. J., Sim, J. H., & Gildersleeve, J. C. (2025). Single-cell glycome and transcriptome profiling enabled by a library of anti-glycan antibodies. Cell Chemical Biology, 32(12), 1554–1568.e1558. 10.1016/j.chembiol.2025.11.005

Nakashima, J., Pattathil, S., Avci, U., Chin, S., Alan Sparks, J., Hahn, M. G., Gilroy, S., & Blancaflor, E. B. (2023). Glycome profiling and immunohistochemistry uncover changes in cell walls of Arabidopsis thaliana roots during spaceflight. npj Microgravity, 9(1). 10.1038/s41526-023-00312-0

Okekeogbu, I. O., Pattathil, S., González Fernández-Niño, S. M., Aryal, U. K., Penning, B. W., Lao, J., Heazlewood, J. L., Hahn, M. G., McCann, M. C., & Carpita, N. C. (2019). Glycome and Proteome Components of Golgi Membranes Are Common between Two Angiosperms with Distinct Cell-Wall Structures. The Plant Cell, 31(5), 1094–1112. 10.1105/tpc.18.00755

Pattathil, S., Avci, U., Baldwin, D., Swennes, A. G., McGill, J. A., Popper, Z., Bootten, T., Albert, A., Davis, R. H., Chennareddy, C., Dong, R., O’Shea, B., Rossi, R., Leoff, C., Freshour, G., Narra, R., O’Neil, M., York, W. S., & Hahn, M. G. (2010). A Comprehensive Toolkit of Plant Cell Wall Glycan-Directed Monoclonal Antibodies. Plant Physiology, 153(2), 514–525. 10.1104/pp.109.151985

Pattathil, S., Avci, U., Miller, J. S., & Hahn, M. G. (2012). Immunological Approaches to Plant Cell Wall and Biomass Characterization: Glycome Profiling. In M. E. Himmel (Ed.), Biomass Conversion: Methods and Protocols (pp. 61–72). Humana Press. 10.1007/978-1-61779-956-3_6

Pattathil, S., Hahn, M. G., Dale, B. E., & Chundawat, S. P. S. (2015). Insights into plant cell wall structure, architecture, and integrity using glycome profiling of native and AFEX^TM^-pre-treated biomass. Journal of Experimental Botany, 66(14), 4279–4294. 10.1093/jxb/erv107

Pierce, S. E., Davis, R. W., Nislow, C., & Giaever, G. (2007). Genome-wide analysis of barcoded Saccharomyces cerevisiae gene-deletion mutants in pooled cultures. Nature Protocols, 2(11), 2958–2974. 10.1038/nprot.2007.427

Puhlmann, J., Bucheli, E., Swain, M. J., Dunning, N., Albersheim, P., Darvill, A. G., & Hahn, M. G. (1994). Generation of Monoclonal Antibodies against Plant Cell-Wall Polysaccharides (I. Characterization of a Monoclonal Antibody to a Terminal [alpha]-(1->2)-Linked Fucosyl-Containing Epitope. Plant Physiology, 104(2), 699–710. 10.1104/pp.104.2.699

Ruprecht, C., Bartetzko, M. P., Senf, D., Dallabernadina, P., Boos, I., Andersen, M. C. F., Kotake, T., Knox, J. P., Hahn, M. G., Clausen, M. H., & Pfrengle, F. (2017). A Synthetic Glycan Microarray Enables Epitope Mapping of Plant Cell Wall Glycan-Directed Antibodies. Plant Physiology, 175(3), 1094–1104. 10.1104/pp.17.00737

Schmidt, D., Schuhmacher, F., Geissner, A., Seeberger, P. H., & Pfrengle, F. (2015). Automated Synthesis of Arabinoxylan-Oligosaccharides Enables Characterization of Antibodies that Recognize Plant Cell Wall Glycans. Chemistry – A European Journal, 21(15), 5709–5713. 10.1002/chem.201500065

Somerville, C., Bauer, S., Brininstool, G., Facette, M., Hamann, T., Milne, J., Osborne, E., Paredez, A., Persson, S., Raab, T., Vorwerk, S., & Youngs, H. (2004). Toward a Systems Approach to Understanding Plant Cell Walls. Science, 306(5705), 2206–2211. 10.1126/science.1102765

Thorne, K., Urbanowicz, B., & Hahn, M. (2024). Plant Cell Wall Glycan-Directed Monoclonal Antibodies. In (pp. 206–236). 10.1201/9781003178309-10

Verhertbruggen, Y., Marcus, S. E., Haeger, A., Ordaz-Ortiz, J. J., & Knox, J. P. (2009). An extended set of monoclonal antibodies to pectic homogalacturonan. Carbohydrate Research, 344(14), 1858–1862. 10.1016/j.carres.2008.11.010

Verhertbruggen, Y., Marcus, S. E., Haeger, A., Verhoef, R., Schols, H. A., McCleary, B. V., McKee, L., Gilbert, H. J., & Paul Knox, J. (2009). Developmental complexity of arabinan polysaccharides and their processing in plant cell walls. The Plant Journal, 59(3), 413–425. 10.1111/j.1365-313x.2009.03876.x

Villalobos, J. A., Yi, B. R., & Wallace, I. S. (2015). 2-Fluoro-L-Fucose Is a Metabolically Incorporated Inhibitor of Plant Cell Wall Polysaccharide Fucosylation. PLoS ONE, 10(9), e0139091. 10.1371/journal.pone.0139091

Walker, J. A., Pattathil, S., Bergeman, L. F., Beebe, E. T., Deng, K., Mirzai, M., Northen, T. R., Hahn, M. G., & Fox, B. G. (2017). Determination of glycoside hydrolase specificities during hydrolysis of plant cell walls using glycome profiling. Biotechnology for Biofuels, 10(1). 10.1186/s13068-017-0703-6

Wilkop, T., Pattathil, S., Ren, G., Davis, D. J., Bao, W., Duan, D., Peralta, A. G., Domozych, D. S., Hahn, M. G., & Drakakaki, G. (2019). A Hybrid Approach Enabling Large-Scale Glycomic Analysis of Post-Golgi Vesicles Reveals a Transport Route for Polysaccharides. The Plant Cell, 31(3), 627–644. 10.1105/tpc.18.00854

Willats, W. G. T., Orfila, C., Limberg, G., Buchholt, H. C., van Alebeek, G.-J. W. M., Voragen, A. G., Marcus, S. E., Christensen, T. M. I. E., Mikkelsen, J. D., Murray, B. S., & Knox, J. P. (2001). Modulation of the Degree and Pattern of Methyl-esterification of Pectic Homogalacturonan in Plant Cell Walls: IMPLICATIONS FOR PECTIN METHYL ESTERASE ACTION, MATRIX PROPERTIES, AND CELL ADHESION*. Journal of Biological Chemistry, 276(22), 19404–19413. 10.1074/jbc.M011242200

Xiao, P., Sahu, P., Pfaff, S. A., Ankur, A., Ranasinghe, Y. K., Gow, N. A. R., Latgé, J.-P., Cosgrove, D. J., & Wang, T. (2025). Revealing structure and shaping priorities in plant and fungal cell wall architecture via solid-state NMR. The Cell Surface, 14, 100159. 10.1016/j.tcsw.2025.100159

Zhong, S., Wang, R., Gao, X., Guo, Q., Lin, R., & Luo, M. (2025). Modular DNA Barcoding of Nanobodies Enables Multiplexed in situ Protein Imaging and High-throughput Biomolecule Detection. eLife Sciences Publications, Ltd. 10.7554/elife.105225.2

